# Synaptic homeostasis at the Drosophila neuromuscular junction is a reversible signaling process that is sensitive to high temperature

**DOI:** 10.1101/154930

**Authors:** Catherine J. Yeates, C. Andrew Frank

## Abstract

Homeostasis is a vital mode of biological self-regulation. The hallmarks of homeostasis for any biological system are a baseline set point of physiological activity, detection of unacceptable deviations from the set point, and effective corrective measures to counteract deviations. Homeostatic synaptic plasticity (HSP) is a form of neuroplasticity in which neurons and circuits resist environmental perturbations in order to maintain appropriate levels of activity. One assumption is that if an environmental perturbation triggers homeostatic corrective changes in neuronal properties, those corrective measures should be reversed upon removal of the perturbation. We test the reversibility and limits of HSP at a well-studied model synapse, the *Drosophila melanogaster* neuromuscular junction (NMJ). At the Drosophila NMJ, impairment of glutamate receptors causes a decrease in quantal size, which is offset by a corrective, homeostatic increase in the number of vesicles released per evoked presynaptic stimulus, or quantal content. This process has been termed presynaptic homeostatic potentiation (PHP). Taking advantage of a GAL4/GAL80^TS^/UAS expression system, we triggered PHP by expressing a dominant-negative glutamate receptor subunit at the NMJ. We then reversed PHP by halting expression of the dominant-negative receptor. Our data show that PHP is fully reversible over a time course of 48-72 hours after the dominant-negative glutamate receptor stops being genetically expressed. Additionally, we found that the PHP response triggered by the dominant-negative subunit was ablated at high temperatures. Our data show that the long-term maintenance of PHP at the Drosophila NMJ is a reversible regulatory process that is sensitive to temperature.

**SIGNIFICANCE STATEMENT:** Biological homeostatic systems must upregulate or downregulate cellular parameters in order to maintain appropriate set points of physiological activity. Homeostasis is a well-documented mode of regulation in metazoan nervous systems. True homeostatic control should be a reversible process – but due to technical difficulties of presenting and removing functional challenges to living synapses, the reversibility of homeostatic forms of synapse regulation has not been rigorously examined *in vivo* over extended periods of developmental time. Here we formally demonstrate that homeostatic regulation of *Drosophila melanogaster* neuromuscular synapse function is reversible and temperature-labile. This is significant because developing methods to study how homeostatic regulatory systems are turned on and off could lead to fundamental new insights about control of synaptic output.

## INTRODUCTION

Homeostasis is a strong form of biological regulation. It permits individual cells or entire systems of cells to maintain core physiological properties that are compatible with life. In the nervous system, decades of study have shown that while synapses and circuits are inherently plastic, they also possess robust homeostatic regulatory systems in order to maintain physiological stability. Homeostatic plasticity in the nervous system is a non-Hebbian strategy to counteract challenges to neuronal function that may threaten to disrupt essential neuronal and circuit activities (Turrigiano, 2017). Depending upon the synaptic preparation examined and the environmental challenge presented to the synapse, homeostatic responses may be executed via compensatory adjustments to presynaptic neurotransmitter release (Cull-Candy et al., 1980; Davis and Müller, 2015; Frank et al., 2006; Murthy et al., 2001; Petersen et al., 1997; Thiagarajan et al., 2005), postsynaptic neurotransmitter receptor composition (O’Brien et al., 1998; Rongo and Kaplan, 1999; Turrigiano, 2008; Turrigiano et al., 1998), neuronal excitability (Bergquist et al., 2010; Marder and Bucher, 2007; Marder and Goaillard, 2006; Marder and Prinz, 2002; Parrish et al., 2014) – or even developmentally, via changes in synaptic contact formation and maintenance (Burrone et al., 2002; Davis and Goodman, 1998; Wefelmeyer et al., 2016).

Bi-directionality has been documented in several homeostatic systems, perhaps most prominently in the case of synaptic scaling of neurotransmitter receptors. For vertebrate neuronal culture preparations – such as cortical neurons or spinal neurons – global silencing of network firing can induce increases in excitatory properties, such as increased AMPA-type glutamate receptor accumulation; by contrast, global enhancement of activity can induce the opposite type of response (O’Brien et al., 1998; Turrigiano, 2008; Turrigiano et al., 1998; Wierenga et al., 2005).

Bi-directionality is also a key feature underlying homeostatic alterations of neurotransmitter release at peripheral synapses like the neuromuscular junction (NMJ). At the *Drosophila melanogaster* and mammalian NMJs, impairing neurotransmitter receptor function postsynaptically results in diminished sensitivity to single vesicles of transmitter. Electrophysiologically, this manifests as decreased quantal size. NMJs respond to this challenge by enhancing neurotransmitter vesicle release (Cull-Candy et al., 1980; Davis et al., 1998; Frank et al., 2009; Petersen et al., 1997; Plomp et al., 1995; Plomp et al., 1992). By contrast, perturbations that enhance quantal size – for example, overexpression of the vesicular neurotransmitter transporter – can result in decreased quantal content (Daniels et al., 2004; Gaviño et al., 2015).

Synapses and circuits possess myriad solutions to assume appropriate functional outputs in the face of perturbations (Marder and Goaillard, 2006; Marder and Prinz, 2002). Therefore, a corollary to bi-directional regulation is that homeostatic forms of regulation should also be reversible. There are experimental difficulties of presenting and removing a synaptic challenge in the context of a living synapse, so homeostatic reversibility has not been rigorously studied in an *in vivo* system or over extended periods of developmental time. However, understanding how homeostatic regulatory systems are reversibly turned on and off could have profound implications for elucidating fundamental properties of circuit regulation.

Here we focus on the aforementioned *Drosophila melanogaster* NMJ as a living synapse to test homeostatic reversibility. At the Drosophila NMJ, a canonical way to challenge synapse function is through glutamate receptor impairment (Frank, 2014), either genetically (Petersen et al., 1997) or pharmacologically (Frank et al., 2006). Impairments of muscle glutamate receptor function decrease quantal size. Decreased quantal size spurs muscle-to-nerve signaling that ultimately results in a homeostatic increase in presynaptic vesicle release – a process that has been termed presynaptic homeostatic potentiation (PHP). The most widely used experimental homeostatic challenges to Drosophila NMJ function are not easily reversed – including pharmacological inhibition with PhTox, due to an irreversible impairment of glutamate receptor function (Frank et al., 2006).

For this study, we engineered a way to challenge NMJ function *in vivo* for significant periods of time, verify the effectiveness of the challenge at a defined time developmental time point, remove the challenge, and then assess the homeostatic capacity of the NMJ at a later developmental time point. To do this, we utilized the TARGET (temporal and regional gene expression targeting) GAL4/GAL80^TS^/UAS expression system (McGuire et al., 2003). By using this expression system to temporally control the expression of a dominant-negative GluRIIA receptor subunit (DiAntonio et al., 1999), we found that homeostatic potentiation of neurotransmitter release is fully reversible. In the course of conducting our studies, we also uncovered a high temperature limitation of homeostatic potentiation at the NMJ.

## MATERIALS AND METHODS

### Drosophila Stocks and Husbandry

Fruit fly stocks were either obtained from the Bloomington Drosophila Stock Center (BDSC, Bloomington, Indiana) or from the labs that generated them. *w^1118^* was used as a wild-type (WT) control (Hazelrigg et al., 1984). The *GluRIIA^SP16^* deletion was used as a genetic loss-of-function (Petersen et al., 1997). Transgenes included the *UAS-*driven dominant-negative glutamate receptor subunit, *UAS-GluRIIA^M614R^* (DiAntonio et al., 1999) and a ubiquitous *Tubulin_Promoter_-Gal80^TS^* (*Tub_P_-Gal80^TS^*) (McGuire et al., 2003). Muscle-specific GAL4 drivers in-cluded *MHC-Gal4* (Schuster et al., 1996a, b) and *BG57-Gal4* (also known as *C57-Gal4*) (Budnik et al., 1996). For reversibility experiments, the full genotypes for the crosses were *w^1118;^ CyO-GFP/UAS-GluRIIA^M614R^; TM6b(Tb)/Tub_P_-Gal80^TS^* x *w^1118^; ; TM6b(Tb)/MHC-Gal4 or BG57-Gal4*. Non-tubby, non-GFP larvae were selected for recording. In control recordings, we found no discernable differences between male and female third-instar electrophysiology, but for reversibility experiments (Figs. 3-5), single sexes of larvae were chosen to eliminate sex as a possible confounding variable.

Fruit flies were raised on cornmeal, molasses, and yeast medium (see BDSC website for standard recipe) in temperature-controlled conditions. For most experiments, animals were reared at the temperatures noted (including temperature shifts) until they reached the wandering third instar larval stage, at which point they were chosen for electrophysiological recording. For experiments in Figs. 2 and 3, mated animals were placed at either 25°C or 29°C and allowed to lay eggs for 6-8 hours. Stage-matched and size-matched early third instar larvae (approximately 48-54 hours after the egg laying period) were subjected to electrophysiological recording (Fig. 2) or temperature swaps (Fig. 3), as indicated.

### Electrophysiology and Analysis

For Figs. 1 and 3-6, wandering third instar larvae were used for electrophysiological recordings. For Fig. 2, early third instar larvae were used. In both cases, sharp electrode electrophysiological recordings were taken from muscle 6 of abdominal segments 2 and 3. Briefly, larvae were dissected in a modified HL3 saline comprised of: NaCl (70 mM), KCl (5 mM), MgCl_2_ (10 mM), NaHCO_3_ (10 mM), sucrose (115 mM = 3.9%), trehalose (4.2 mM = 0.16%), HEPES (5.0 mM = 0.12%), and CaCl_2_ (0.5 mM). Electrophysiological data were collected using an Axopatch 200B amplifier (Molecular Devices, Sunnyvale, CA) in bridge mode, digitized using a Digidata 1440A data acquisition system (Molecular Devices), and recorded with pCLAMP 10 acquisition software (Molecular Devices). A Master-8 pulse stimulator (A.M.P. Instruments, Je-rusalem, Israel) and an ISO-Flex isolation unit (A.M.P. Instruments) were utilized to deliver 1 ms suprathreshold stimuli to the appropriate segmental nerve. The average spontaneous miniature excitatory postsynaptic potential (mEPSP) amplitude per NMJ was quantified by hand, approximately 100-200 individual spontaneous release events per NMJ (MiniAnalysis, Synaptosoft, Fort Lee, NJ). In the case that the mEPSP frequency was extremely low (usually for expression of the dominant-negative glutamate receptor subunit), several minutes of spontaneous recording were done and all events measured. Measurements from all NMJs of a given condition were then averaged. For evoked neurotransmission, 30 excitatory postsynaptic potentials (EPSPs) were averaged to find a value for each NMJ. These were then averaged to calculate a value for each condition. Quantal content (QC) was calculated by the ratio of average EPSP and average mEPSP amplitudes for each individual NMJ. An average quantal content was then calculated for each condition.

**Figure 1.**
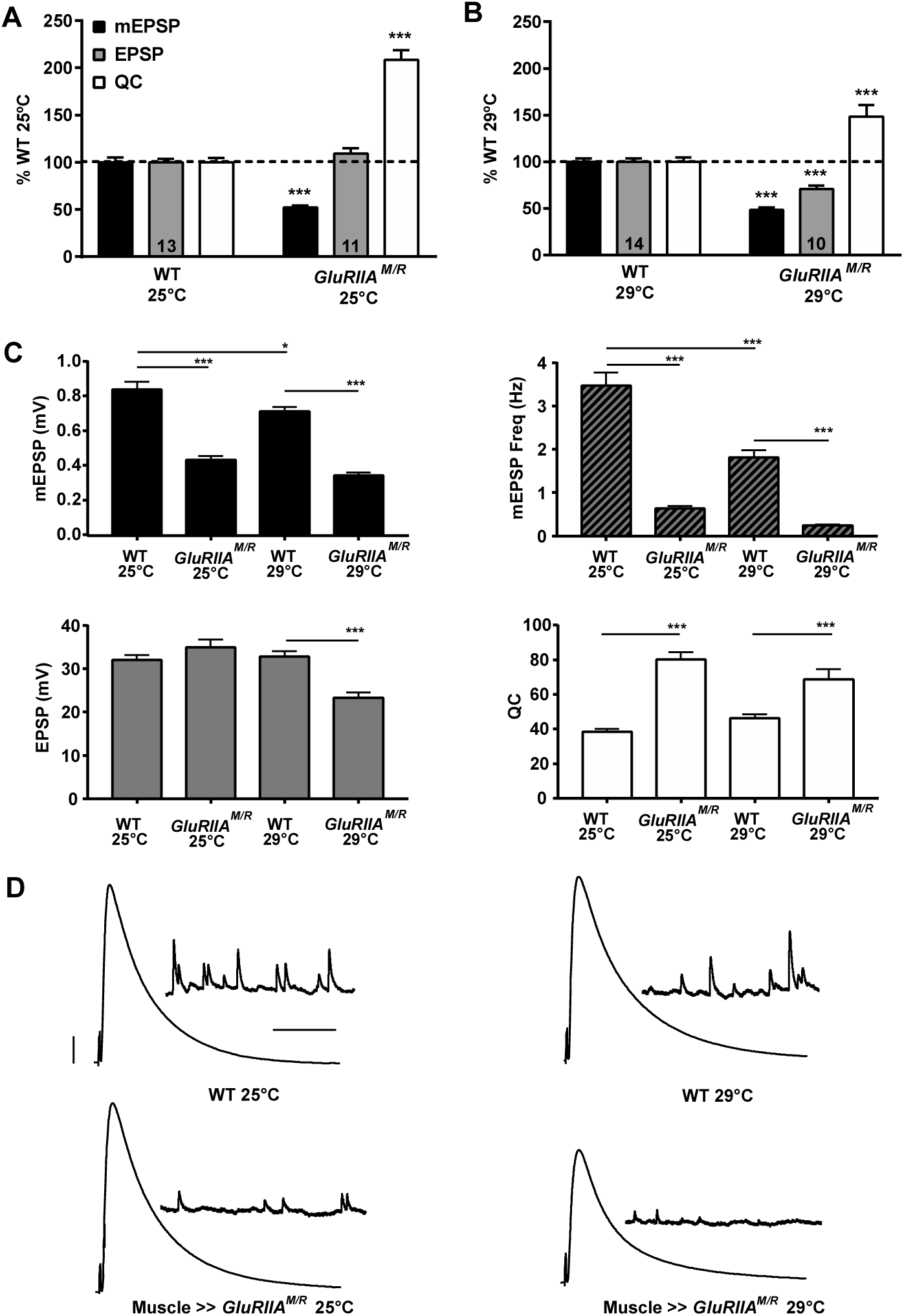
Postsynaptic expression of GluRIIA^M/R^ causes a decrease in mEPSP size and a homeostatic increase in quantal content. **(A)** Electrophysiological profiles comparing *w1118* (WT) control NMJs to NMJs with postsynaptic expression of *UAS-GluRIIA^M/R^* (*w; UASGluRIIA^M/R^/+; MHC-Gal4/+*). Animals were reared at 25°C. mEPSP amplitude is markedly decreased compared to WT (*** *p* < 0.001). Quantal content (QC) is significantly increased for the dominant-negative NMJs (*** *p* < 0.001), showing a homeostatic response that maintains EPSP amplitude at control levels. **(B)** The same genotypes were raised at 29°C. mEPSP amplitude is significantly decreased in the dominant-negative (*** *p* < 0.001). EPSP size is also decreased in the dominant-negative compared to WT (*** *p* < 0.001), with QC significantly increased (*** *p* < 0.001) but not enough to completely offset the decrease in quantal size. **(C)** Raw value comparisons of the recordings normalized in (A) and (B). mEPSP amplitude, mEPSP frequency, EPSP amplitude and QC were compared for these four conditions: WT 25°C, WT 29°C, *w; GluRIIA^M/R/^+; MHC-GAL4/+* 25°C, and *w; GluRIIA^M/R/^+; MHC-GAL4/+* 29°C. In addition to the observations above, WT at 29°C had a decreased mEPSP amplitude (* *p* <0.05) and mEPSP frequency (*** *p* < 0.001) compared to WT at 25°C. There was no difference in quantal content between the two WT conditions (*p* = 0.32). At 25°C, the dominant-negative showed a decrease in mEPSP frequency compared to WT at the same temperature (*** *p* < 0.001). The dominant-negative at 29°C also showed a significant decrease in mEPSP frequency compared to WT at 29°C (*** *p* < 0.001). **(D)** Electrophysiological traces. Scale bars for EPSPs (and mEPSPs) are 5 mV (1 mV) and 50 ms (1000 ms). All statistical comparisons done by one-way ANOVA with Tukey’s post-hoc, collectively comparing the four total conditions.

**Figure 2.**
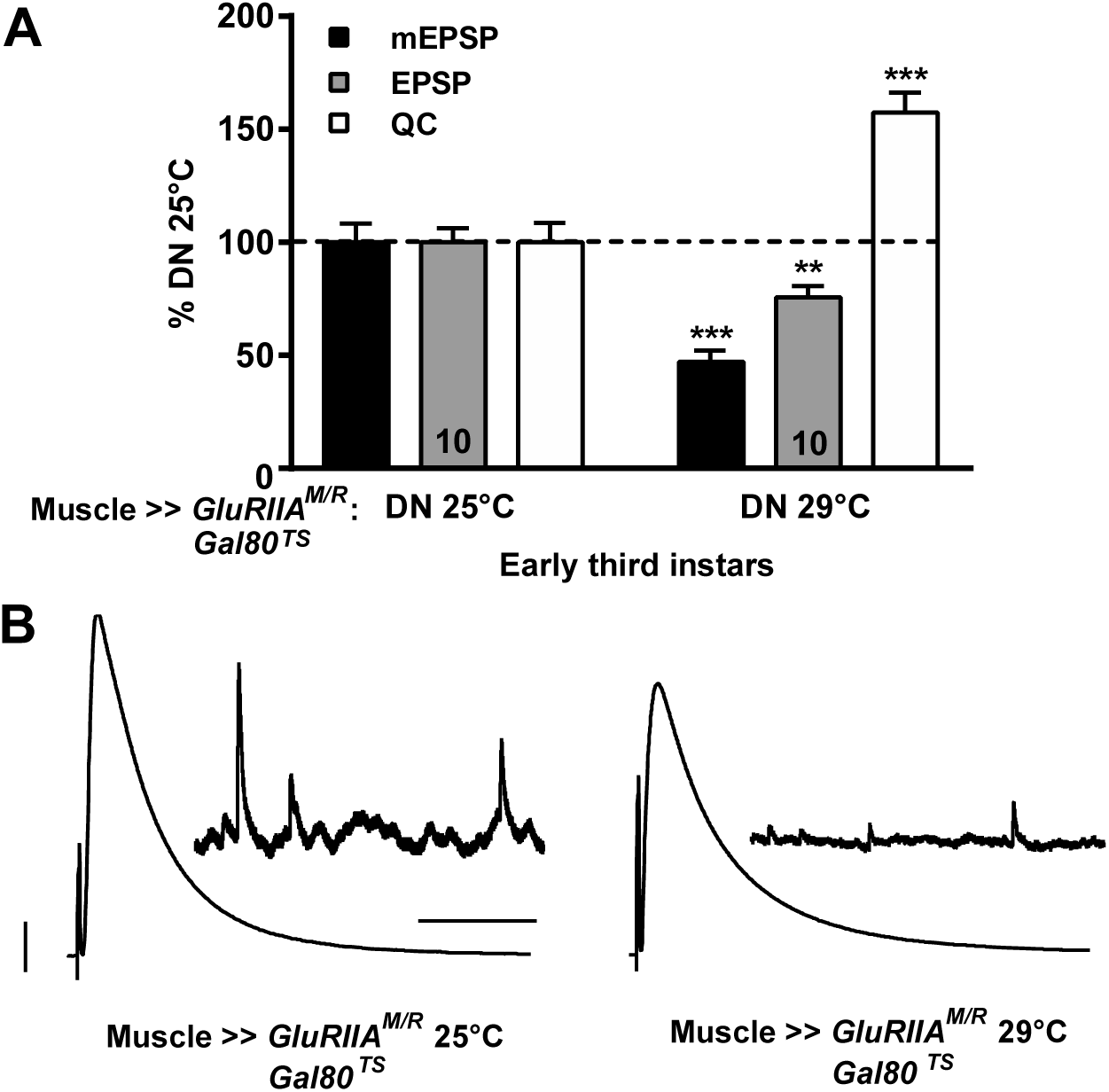
Animals expressing GluRIIA^M/R^ show homeostatic compensation early in development. **(A)** *w; GluRIIA^M/R^/+; MHC-GAL4/GAL80^TS^* animals were reared at 25°C or 29°C. Compared to animals reared at 25°C, the animals reared at 29°C had a decrease in mEPSP amplitude (*** *p* < 0.001, Student’s T-test) and frequency (*p* <0.001, see Table 1). EPSP amplitude was decreased (** *p* < 0.01), but there was a significant increase in QC (*** *p* < 0.001) **(B)** Representative electrophysiological traces. Scale bars for EPSPs (and mEPSPs) are 5 mV (1 mV) and 50 ms (1000 ms).

**Figure 3.**
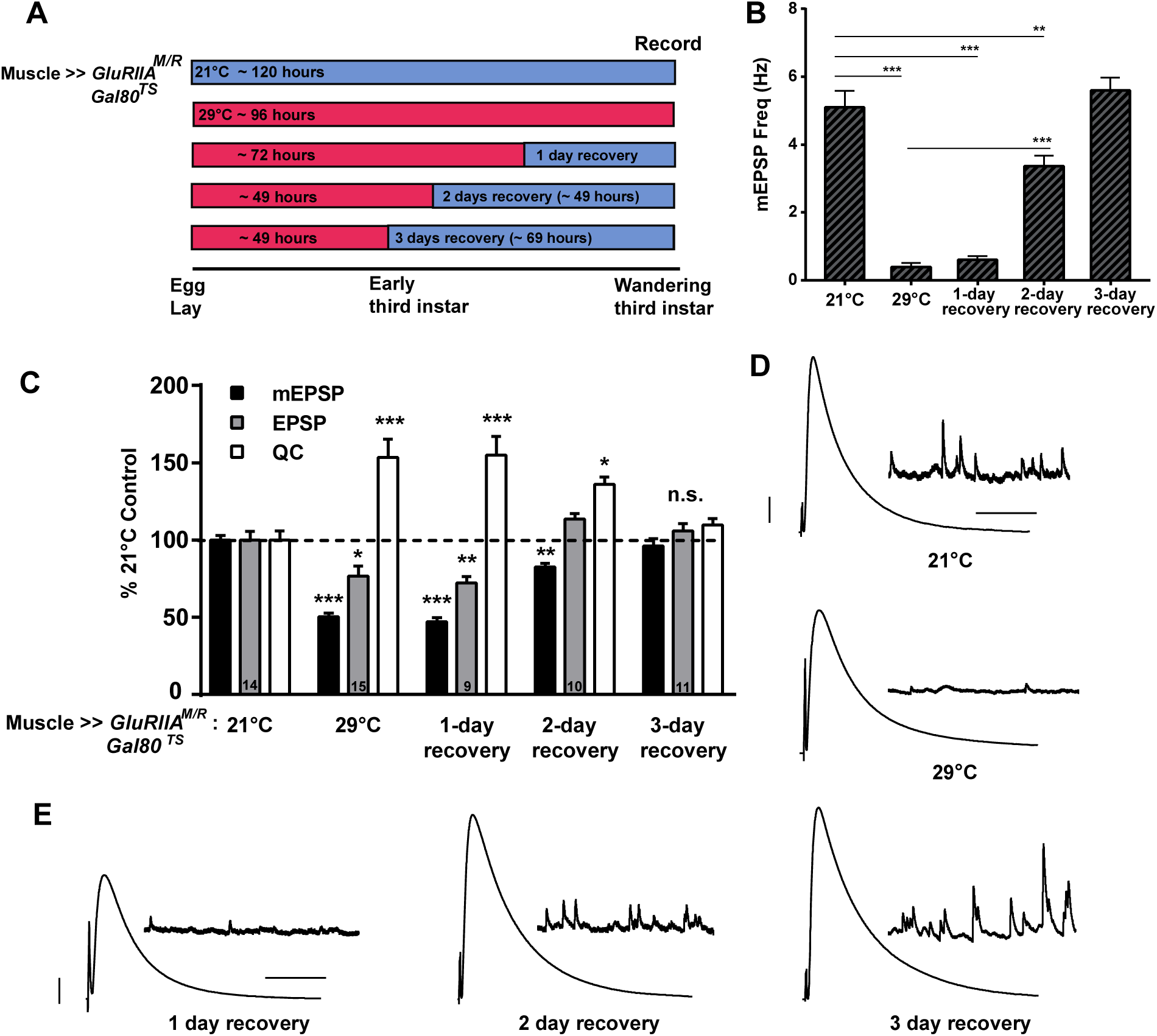
Presynaptic homeostatic potentiation is reversible. **(A)** Diagram of a temperature swap paradigm. Mated animals were allowed to lay eggs for 6-8 hours. One set of animals was reared entirely at 21° from egg lay to electrophysiological recording. A second condition was raised entirely at 29°C. To test for reversibility of homeostatic potentiation, animals were reared initially at 29°C and then swapped to 21°C. Animals were allowed to recover for either 1, 2, or 3 days before recording. **(B)** Expression of the dominant-negative transgene throughout life (29°C) causes a dramatic decrease in mEPSP frequency (*** *p* < 0.001, one-way ANOVA with Tukey’s post-hoc). Frequency remains low after 1 day of recovery at 21°C, compared 21°C rearing controls. By 2 days recovery, the frequency is significantly increased compared to 29°C (*** *p* < 0.001), but still significantly decreased compared to 21°C controls (** *p* < 0.01). By 3 days recovery, mEPSP frequency is no different from animals raised entirely at 21°C. **(C)** Normalized electrophysiological data for mEPSP amplitude, EPSP amplitude, and QC for the NMJs of animals raised as described in (A). Longer recovery periods yield electrophysiology that more closely approximates the control 21°C rearing condition (* *p* < 0.05, ** *p* < 0.01, *** *p* < 0.001 compared to 21°C control by one-way ANOVA with Tukey’s post-hoc). **(D)** Representative electrophysiological traces for 21°C and 29°C. Scale bars refer to 5 mV and 50 ms for EPSPs and 1 mV and 1000 ms for mEPSPs. **(E)** Traces for recovery conditions. Scale bars for EPSPs (and mEPSPs) are 5 mV (1 mV) and 50 ms (1000 ms) for both (D) and (E).

### Statistical Analyses

Statistical analyses were conducted using GraphPad Prism Software. Statistical significance was assessed either by Student’s T-Test when one experimental data set was being directly compared to a control data set, or one-way ANOVA with Tukey’s post-hoc test when multiple data sets were being compared. To assess potential correlations between incubation temperature recovery times and electrophysiological parameters (Fig. 4), Pearson correlation coefficients were calculated (r) and reported on the graphs, and two-tailed statistical analyses performed to check correlation significance.

**Figure 4.**
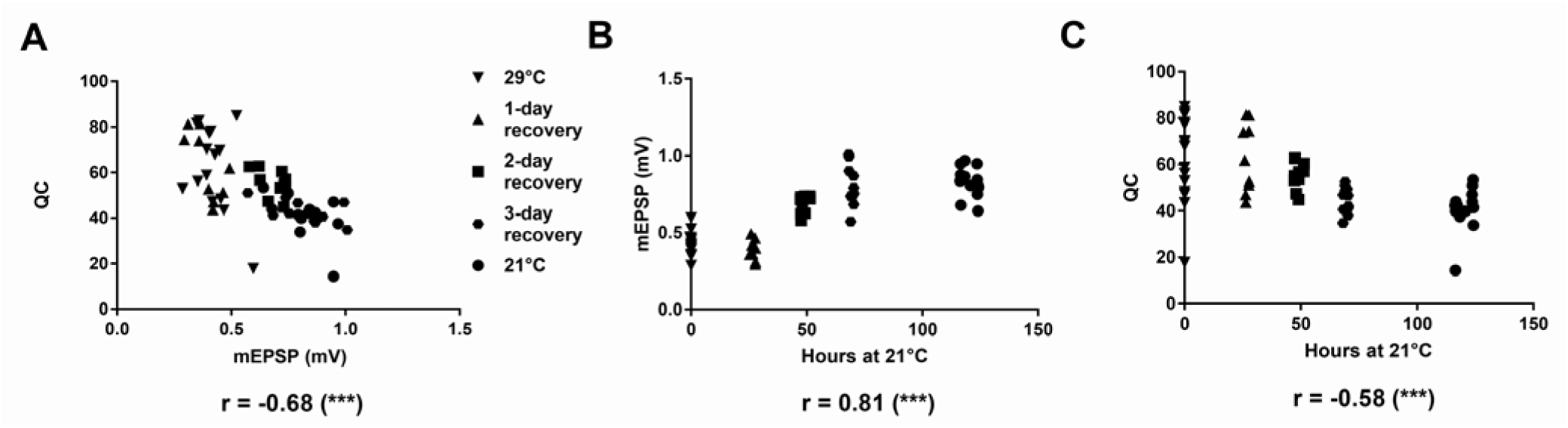
Strong correlations between recovery time and classical homeostatic parameters. **(A)** There is a significant inverse correlation between mEPSP amplitude and QC for the swap experiments described in Figure 3 (Pearson’s *r* = -0.68, *** *p* < 0.001, *n* = 59). **(B)** There is a significant positive correlation between the number of hours an animal has spent at 21°C and mEPSP amplitude. (*r* = 0.81, *p*<0.001, *n* = 59). Animals reared for “0 hours” at 21°C are the 29°C condition, with animals in the 21°C condition being reared at that temperature for around 120 hours before recording. **(C)** A significant inverse correlation between hours at 21°C and quantal content was found (*r* = -0.58, *** *p* < 0.001, *n* = 59). Animals with the highest quantal content were either at 21°C for 0 hours (29°C condition) or 24 hours (1-day recovery).

Specific *p* value ranges are noted in the Fig. legends and shown in graphs as follows: * *p* < 0.05, ** *p* < 0.01, and *** *p* < 0.001. For some *p* values that potentially trend toward statistical significance (0.05 < *p* < 0.1), specific values are given. Most values reported in Table 1 or plotted on bar graphs are mean ± SEM.

**Table 1.**
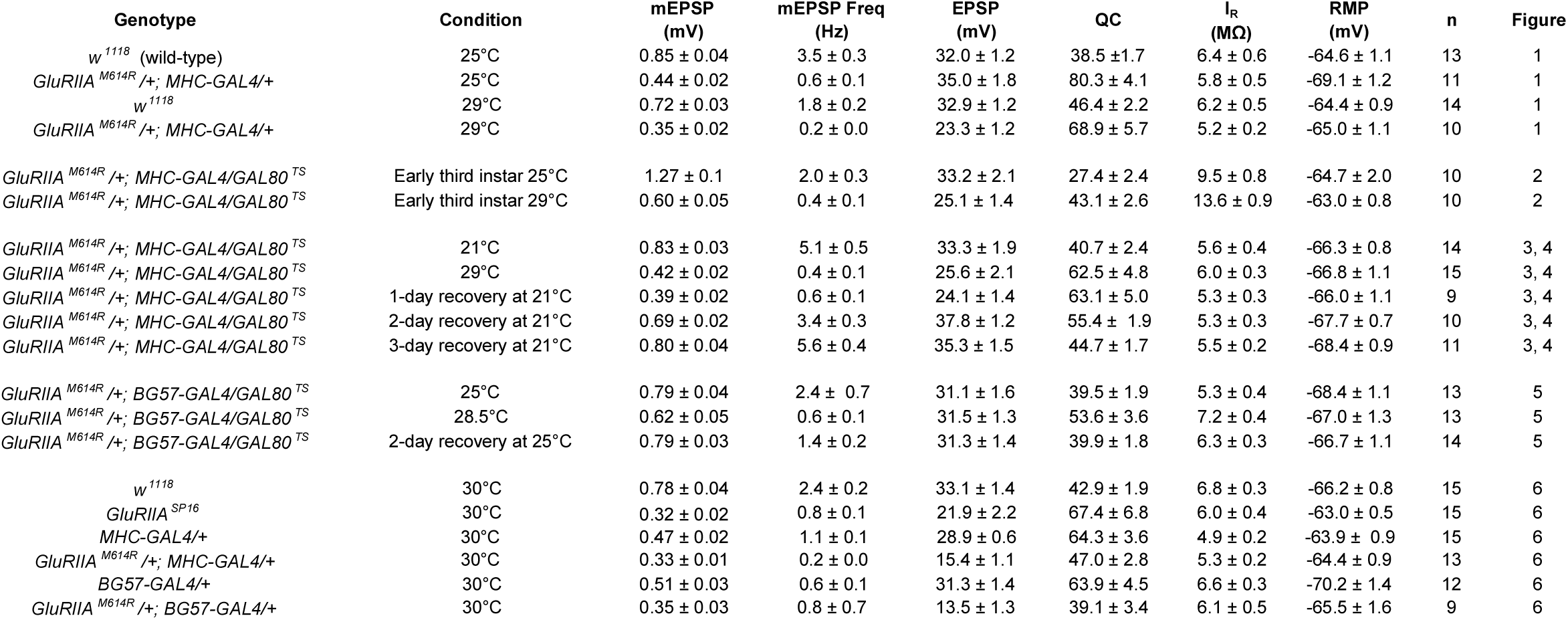
Raw Electrophysiological Data. Full genotypes and rearing conditions for electrophysiological data presented in the study. Average values ± SEM are presented for each parameter, with *n* = number of NMJs recorded. Values include miniature excitatory postsynaptic potential (mEPSP) amplitude, mEPSP frequency, excitatory postsynaptic potential (EPSP) amplitude, quantal content (QC), muscle input resistance, and resting membrane potential.

## RESULTS

### Homeostatic Potentiation Utilizing a Dominant-Negative Glutamate Receptor Subunit

A prior study described a transgene encoding a dominant-negative GluRIIA subunit, *UAS-GluRIIA^M614R^* (herein termed *UAS-GluRIIA^M/R^* or “dominant-negative”) (DiAntonio et al., 1999). The GluRIIA M614R amino-acid substitution resides in the ion conduction pore of the GluRIIA subunit, and it cripples channel function (DiAntonio et al., 1999). Transgenic expression of *UAS-GluRIIA^M/R^* in muscles renders a strong homeostatic challenge (markedly diminished quantal size) and an equally strong compensatory response (increase in quantal content). The prior report demonstrated that evoked amplitudes remain normal as a result of this compensation (DiAntonio et al., 1999).

We acquired this dominant-negative transgenic line to test homeostatic reversibility. First, we replicated the published experiments – this time raising fruit fly larvae at temperatures compatible with the TARGET system that we planned to use to test reversibility (McGuire et al., 2003). We tested 25°C (a common culturing temperature) and 29°C (a temperature at which the GAL80^TS^ protein ceases to inhibit GAL4). We drove *UAS-GluRIIA^M/R^* expression with *MHC-Gal4*, which turns on in first-instar larval muscles (Schuster et al., 1996a, b). We recorded from NMJs of control and dominant-negative wandering third instar larvae.

For animals reared at 25°C, wild-type (WT) electrophysiology was robust (Figs. 1A, C, D, Table 1 – see Table 1 for all raw electrophysiological data). By contrast, muscle-specific *UAS-GluRIIA^M/R^* expression at 25°C caused a large diminishment of quantal size compared to WT, evident from a small average miniature excitatory postsynaptic potential (mEPSP) amplitude (Figs. 1A, C, Table 1). Despite this decrease in quantal size, there was no diminishment in the average evoked excitatory postsynaptic potential (EPSP) amplitude for dominant-negative NMJs (Figs. 1A, C, D, Table 1) because of an offsetting homeostatic increase in quantal content (QC) (Figs. 1A, C, Table 1). In sum, recordings at 25°C agreed with the prior finding for the dominant-negative transgene (DiAntonio et al., 1999): perfect presynaptic homeostatic potentiation (PHP). We did note one previously unreported phenotype at 25°C: starkly diminished quantal frequency (Fig. 1C). This diminished mEPSP frequency phenotype mirrored numerous cases in which there was greatly diminished glutamate receptor subunit expression or function (Brusich et al., 2015; Daniels et al., 2006; Featherstone et al., 2005; Kim et al., 2012; Kim et al., 2015; Ramos et al., 2015).

For animals raised at 29°C, we garnered similar results as 25°C, noting a few differences. First, at 29°C, WT NMJs had slightly smaller mEPSP quantal amplitude and frequency than 25°C WT NMJs (Figs. 1A-D, Table 1). This finding was consistent with a prior report measuring NMJ physiology at elevated temperatures (Ueda and Wu, 2015). Nevertheless, WT evoked amplitudes and QC at 29°C were robust, similar to the values garnered at 25°C (Figs. 1A-D, Table 1). *UAS-GluRIIA^M/R^*-expressing NMJs from 29°C showed a profound decrease in mEPSP size, a strong compensatory increase in QC, and markedly reduced quantal frequency compared to WT controls (Fig. 1B). However, unlike at 25°C, homeostatic compensation was not perfect at 29°C for dominant-negative NMJs. Even though QC was significantly increased in dominant-negative NMJs compared to WT, it did not increase enough to bring average *MHCGal4 >> UAS-GluRIIA^M/R^* EPSP amplitudes at 29°C fully back to control levels (Figs. 1B, C, D).

### Early Third Instar Larvae Express Homeostatic Plasticity

Our initial experiments showed that it is possible to observe compensatory increases in quantal content for *UAS-GluRIIA^M/R^*-expressing animals raised throughout life, at either 25°C or 29°C. We sought to test homeostatic reversibility using the TARGET system. For this study, TARGET augments GAL4/UAS expression adding a ubiquitously expressed *Tubulin_P_*-*Gal80^TS^* GAL4 inhibitor transgene (McGuire et al., 2003) to the *MHC-Gal4* >> *UAS-GluRIIA^M/R^* genetic background. The temperature sensitivity of the GAL80^TS^ protein permits tight control over when GAL4-responsive transgenes are expressed (McGuire et al., 2003). We predicted that the dominant-negative transgene should be repressed by GAL80^TS^ at low, permissive temperatures; and conversely, *UAS-GluRIIA^M/R^* should be actively expressed when GAL80^TS^ is inactive (~29°C or higher) (McGuire et al., 2003). In order to study the reversibility of PHP, animals could be reared at high temperature and then swapped to a lower temperature at an appropriate developmental time point.

We needed to identify a suitable developmental stage for temperature swaps. An ideal swap point would be late enough to detect PHP, but early enough to allow recovery time after cessation of the dominant-negative *UAS-GluRIIA^M/R^* expression. We crossed *UAS-GluRIIA^M/R^*; *TubP-Gal80^TS^* x *MHC-Gal4* stocks. Mated animals were transferred to 25°C or 29°C for egg laying and subsequent larval development. We selected early third instar progeny for electrophysiological recording. At temperature ranges of 25-29°C, early third instar larvae emerge roughly 48-60 hours after an egg-laying period. We staged small animals unambiguously by examining their posterior spiracles for an orange-colored tip.

For animals raised entirely at 25°C, early third instar NMJ mEPSP size was large – significantly larger than one would observe for third instar larvae (Figs 2A, B, Table 1). This was expected because for developing NMJs, small muscles have a significantly greater input resistance and enhanced quantal size (Davis and Bezprozvanny, 2001; Lnenicka and Mellon, 1983a, b) (Table 1). Early third instar larval NMJs showed normal evoked amplitudes, consistent with a stable level of evoked muscle excitation throughout development (Figs 2A, B, Table 1).

We predicted genotypically equivalent early third instars raised entirely at 29°C would express the dominant-negative transgene. As expected, NMJs from these animals showed sharply reduced mEPSP amplitude and frequency compared to their stage- and size-matched counterparts raised and 25°C (Figs. 2A, B, Table 1). There was a robust increase in quantal content at 29°C, resulting in EPSP amplitudes that were nearly normal, but not quite at the same level as at 25°C. As with the earlier experiments, presynaptic homeostatic potentiation (PHP) for early third instar NMJs raised exclusively at 29°C was present, but not perfect (Figs. 2A, B, Table 1).

### Imperfect Homeostatic Potentiation is Reversible

At 29°C, PHP was not perfect, but QC increases versus controls were robust, making it possible to test reversibility. We generated additional *MHC-Gal4* >> *UAS-GluRIIA^M/R^* larvae with the *Tub_P_-GAL80^TS^* transgene. This time we chose 21°C as a permissive GAL80^TS^ shift temperature to examine because 21°C permitted multiple electrophysiological time point measurements over a long recovery window (Fig. 3A).

Control (no PHP) animals raised entirely at 21°C took ~120 hours after the egg laying period to reach the wandering third instar stage, while control (PHP) animals raised entirely at 29°C took ~96 hours to reach the same stage (Fig. 3A). A 1-day recovery condition was utilized – exposing animals to the *UAS-GluRIIA^M/R^* challenge for ~72 hours (29°C) and allowing them to recover at 21°C for 1 day prior to recording. We also tested 2- and 3-day recovery conditions, rearing larvae at 29°C for approximately ~49-50 hours and then swapping them to 21°C until they reached wandering third instar stage. Due to the length of the egg lays and small variations in developmental time, some animals reached wandering third instar stage after about 2 days at 21°C, while others took closer to 3 days (Fig. 3A).

Control *MHC-Gal4* >> *UAS-GluRIIA^M/R^* larvae with ubiquitously expressed *Tubp-Gal80^TS^* and raised at 21°C showed physiology indistinguishable from WT control animals reared at 25°C. This was true for all electrophysiological parameters (Compare Figs. 1, 3, Table 1). By contrast, genetically identical control animals raised at 29°C throughout life showed electrophysiological phenotypes similar to dominant-negative *MHC-Gal4* >> *UAS-GluRIIA^M/R^* animals raised at 29°C (Compare Figs. 1, 3, Table 1). Compared to counterparts raised at 21°C, animals raised at 29°C showed reduced mEPSP frequency, reduced mEPSP size, slightly below-normal EPSP amplitudes, and increased QC, indicating robust PHP (Fig. 3, Table 1).

Recovery conditions showed physiological signatures that corresponded with how much time was spent at 21°C. Compared to the 29°C condition, the 1-day 21°C recovery condition showed no significant changes in physiological properties and no reversal of PHP (Figs 3B-D). By contrast, the 2-day recovery condition showed intermediate phenotypes. Compared to constant exposure to 21°C, the 2-day recovery condition still had diminished mEPSP amplitude and frequency – but not nearly as diminished as constant exposure to 29°C (Figs. 3B-D). Interestingly, the 2-day recovery EPSP amplitudes revealed restored levels of excitation, due to an increase in QC (Figs. 3C, D). The 3-day recovery condition showed electrophysiology that was not significantly different from the constant 21°C condition (Figs 3B-D), indicating a full reversal of PHP.

We further analyzed the aggregate data from the reversibility experiment. We wished to test for hallmarks of PHP and reversal related to recovery time. Prior studies of homeostatic plasticity at the Drosophila NMJ have shown that by plotting hundreds individual recording values, quantal content inversely scales with quantal size across genotypes – and as a result, evoked excitation levels remain stable (Davis and Müller, 2015; Frank et al., 2006; Gaviño et al., 2015). For our temperature swap experiments – this time conducting a comparison within a single genotype – this also proved to be the case (Fig. 4A).

Next, we plotted mEPSP and QC values versus the number of animals spent at the recovery temperature, with a specific recovery time value for each NMJ, time locked to the egg laying period and the recording time after recovery/development at 21°C. Both plots showed hallmarks of PHP reversal: mEPSP amplitudes significantly positively correlated with time at 21°C (Fig. 4B), and QC values significantly inversely correlated with time at 21°C (Fig. 4C). Consistent with the observation that PHP was present – but not perfect – at 29°C (Figs. 1-3), individual data points from NMJs of animals raised entirely at 29°C showed a wide variability of QC values (Fig. 4C).

### Perfect Homeostatic Potentiation is Also Reversible

Homeostatic potentiation was robust, yet imperfect, at the 29°C condition. We hoped to test a condition that was both compatible with perfect PHP and the reversibility assay. We attempted a new temperature swap, changing a few parameters. For one change, we lowered the restrictive Gal80^TS^ temperature from 29°C to 28.5°C. Other studies using the TARGET system in *Drosophila melanogaster* have reported that 28.5°C is somewhat effective at impairing Gal80^TS^ function (Corrigan et al., 2014; Redhai et al., 2016; Staley and Irvine, 2010). Second, there was a formal possibility that imperfect compensation at 29°C reflected a GAL4 driver-specific phenomenon, rather than a temperature-specific phenomenon. Therefore, we replaced the *MHC-Gal4* muscle driver with the *BG57-Gal4* muscle driver (Budnik et al., 1996). Finally, we returned to 25°C as the permissive condition.

We generated new sets of larvae for this swap experiment – *BG57-Gal4* >> *UAS-GluRIIA^M/R^* larvae with the *TubP-Gal80^TS^* transgene. As expected, animals raised at 25°C throughout life (GAL80^TS^ on) developed NMJs with electrophysiological properties similar to other control conditions already reported (Figs. 5A, C, Table 1). Also, as expected, animals raised at 28.5°C throughout life (GAL80^TS^ impaired) had significantly diminished NMJ mEPSP amplitudes and mEPSP frequency (Figs. 5A, B, Table 1). NMJs from those 28.5°C animals also showed completely normal NMJ EPSP amplitudes because of a perfect, offsetting homeostatic increase in QC (Figs. 5A, C, Table 1). Of note, the 28.5°C NMJs had markedly diminished mEPSP frequency, an indication of successful expression of the dominant-negative *GluRIIA^M/R^* transgene (Fig. 1). Animals raised at 28.5°C until early third instar and then swapped to 25°C for the final two days of larval development showed NMJ electrophysiology indistinguishable from that of animals raised at 25°C throughout life, indicating a complete reversal of PHP (Fig. 5).

**Figure 5.**
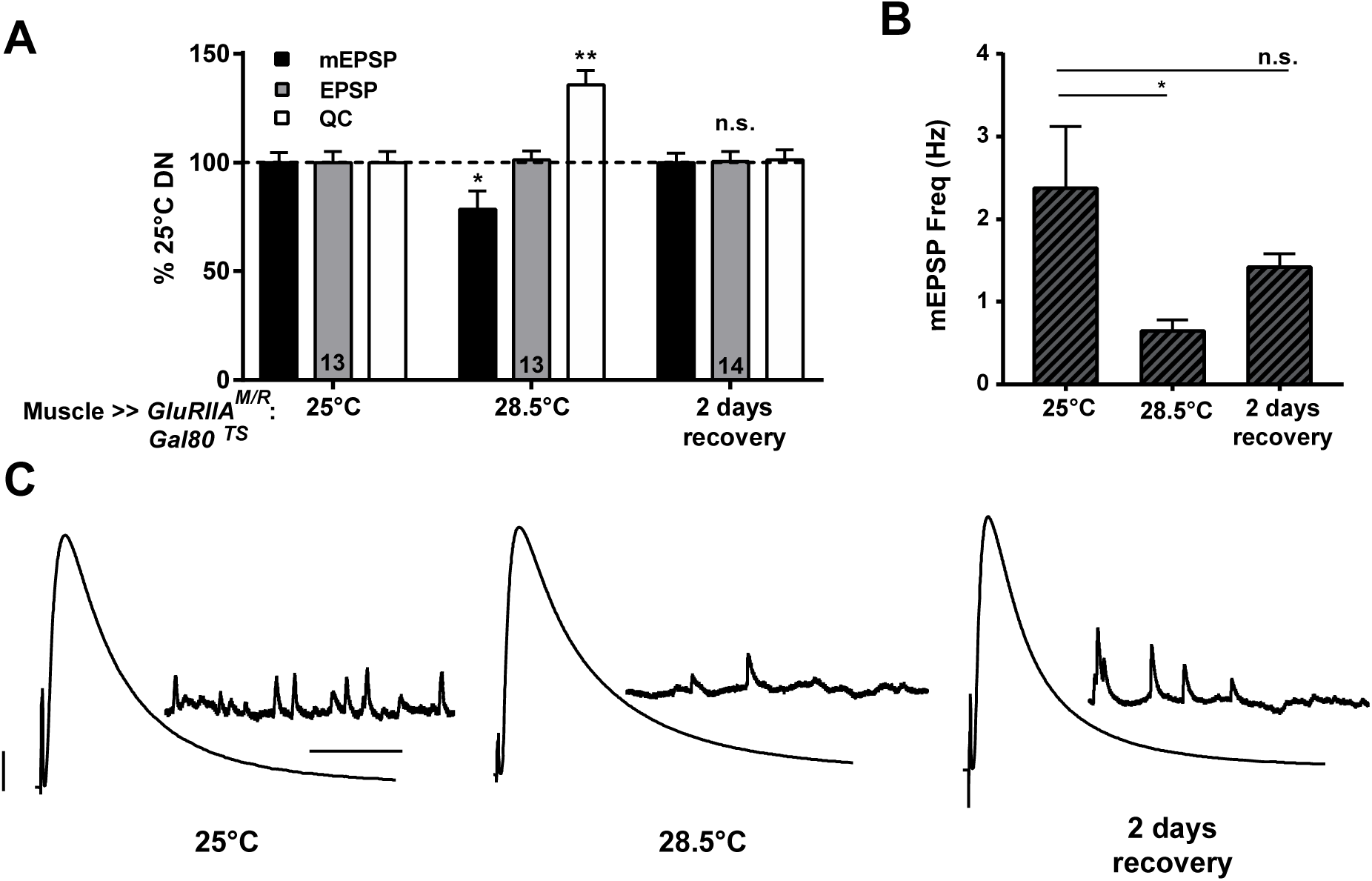
Perfect homeostatic compensation is also reversible. **(A)** Dominant-negative animals raised at 28.5°C show a significant decrease in mEPSP amplitude (* *p* < 0.05, compared to 25°C condition; one-way ANOVA with Tukey’s post-hoc), accompanied by a perfectly offsetting increase in QC (** *p* < 0.01). Reversibility of homeostatic potentiation was demonstrated by initially rearing *w; GluRIIA^M/R^/+; BG57-GAL4/GAL80^TS^* animals at 28.5°C for 2 days and swapping them to 25°C for 2 days. mEPSP amplitude, EPSP amplitude, and quantal content returned to control (25°C throughout life) levels. EPSP amplitudes were virtually identical across all of the conditions. **(B)** Dominant-negative animals raised at 28.5°C show a significant decrease in mEPSP frequency (\**p* < 0.05, one-way ANOVA with Tukey’s post-hoc). By contrast, the mEPSP frequency in animals after two days of recovery was not significantly different from that of the animals reared at 25°C. **(C)** Representative electrophysiological traces. Scale bars for EPSPs (and mEPSPs) are 5 mV (1 mV) and 50 ms (1000 ms).

### Homeostatic potentiation induced by GluRIIA^M/R^ is impaired at high temperatures

*GluRIIA^M/R^* transgene-induced homeostatic potentiation was robust, yet imperfect at the 29°C condition. We wondered if PHP might specifically be impaired at the NMJ when flies are raised at high temperatures. To test this idea further, we drove the dominant-negative transgene in the muscle, setting up crosses to generate both *MHC-Gal4* >> *UAS-GluRIIA^M/R^* and *BG57- Gal4* >> *UAS-GluRIIA^M/R^* animals, as well as driver-specific controls. For the driver controls, quantal size was somewhat diminished at 30°C (Figs. 6A-D, Table 1). This was consistent with the idea that quantal size is generally diminished at very high temperatures (Ueda and Wu, 2015) – though it could also be the case that high levels of muscle-driven GAL4 protein at high temperatures contributes to this phenotype. Evoked EPSP amplitudes were still robust for driver controls (Figs. 6A-D, Table 1). For the dominant-negative NMJs, there was a significant reduction in mEPSP size – beyond what was measured for the driver controls. Moreover, evoked amplitudes were weak, diminished significantly versus driver controls because of no significant increase in QC – in fact, there was a significant decrease in QC (Figs. 6A-D, Table 1). These data indicated that signaling processes that maintain PHP in response to the dominant-negative transgene may break down at high temperatures.

**Figure 6.**
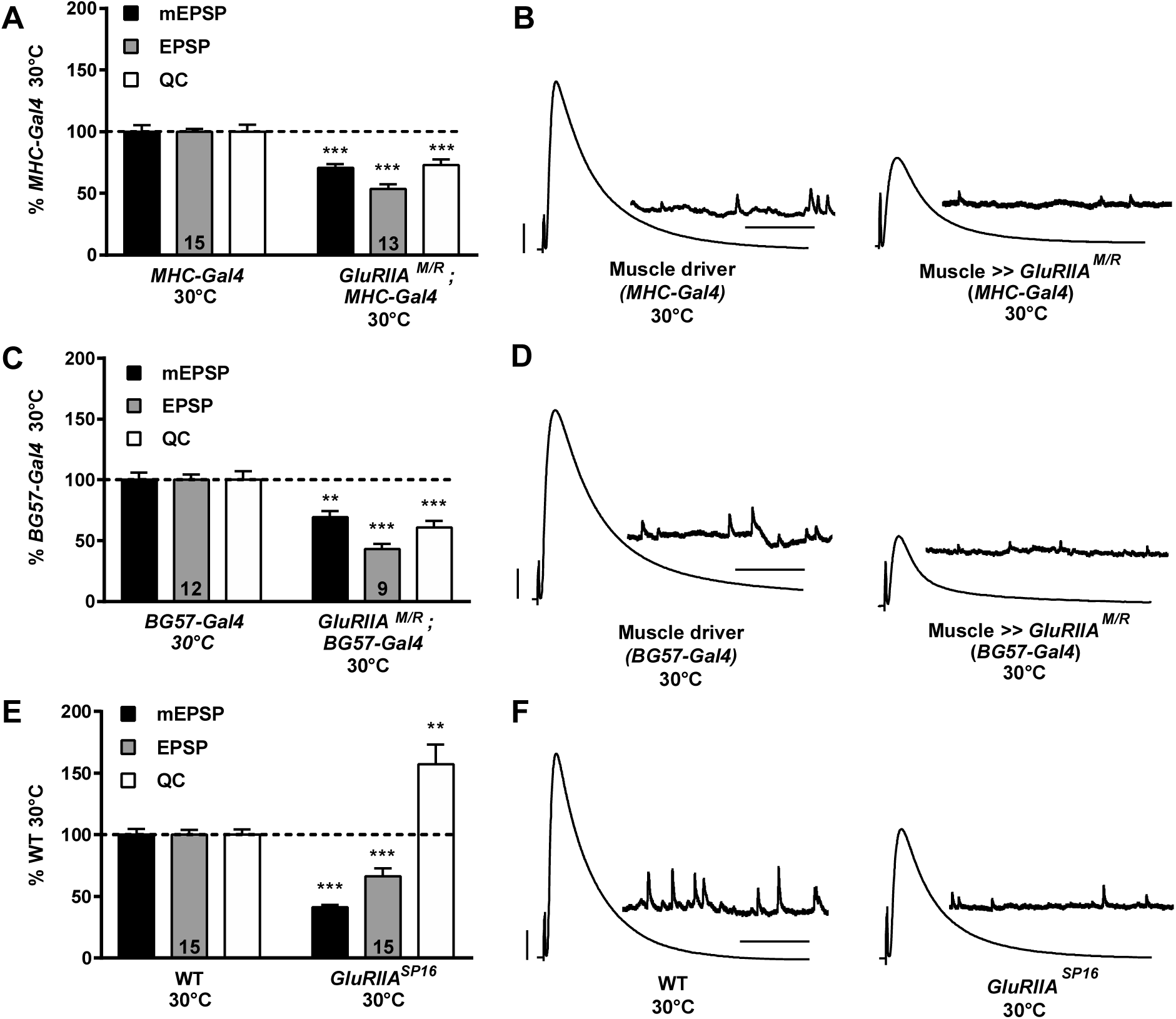
Presynaptic homeostatic potentiation can be impaired at high temperatures. **(A)** *MHC-Gal4* was used to drive expression of the dominant-negative GluRIIA^M/R^ subunit. Dominant-negative animals were compared at 30°C to driver controls, *MHC-GAL4* crossed to WT. mEPSP amplitude was significantly decreased in the dominant-negative animals (*** *p* < 0.001, Student’s T-test). A dramatic decrease in EPSP amplitude also occurred (*** *p*< 0.001) because of a significant decrease in QC in the dominant-negative animals (*** *p* < 0.001). **(B)** Representative electrophysiological traces. Scale bars for EPSPs (and mEPSPs) are 5 mV (1 mV) and 50 ms (1000 ms). **(C)** *BG57-Gal4* was used to drive expression of the dominant-negative GluRIIA^M/R^ subunit. Dominant-negative animals were compared at 30°C to driver controls, *BG57-Gal4* crossed to WT. mEPSP amplitude was significantly decreased for the dominant-negative NMJs (** *p* <0.01, Student’s T-test). A dramatic decrease in EPSP amplitude also occurred (*** *p*<0.001) because of a significant decrease in QC in the dominant-negative animals (*** *p* < 0.001). **(D)** Representative electrophysiological traces. Scale bars as in (B). **(E)** *GluRIIA^SP16^* animals and WT controls were raised at 30°C. mEPSPs were significantly decreased for *GluRIIA* mutants (Student’s T-test, *** *p* < 0.001, Student’s T-test) and QC was significantly increased (** *p* < 0.01). EPSP values were also decreased (*** *p* < 0.001) but not to the same degree as mEPSP amplitudes. **(F)** Representative electrophysiological traces. Scale bars as in (B).

To extend this line of inquiry, we examined a second homeostatic challenge to NMJ function. We raised WT and *GluRIIA^SP16^* deletion flies (Petersen et al., 1997) at 30°C. WT NMJs showed relatively normal physiology at 30°C (Figs., 6E, F, Table 1). NMJs from *GluRIIA^SP16^* deletion animals raised at 30°C showed significant PHP, with reduced mEPSP amplitudes and robust increase in QC compared to WT (Figs. 6A, B, Table 1). We noted that EPSP amplitudes were somewhat reduced compared to WT (Table 1). Collectively, our data show that normal baseline neurotransmission is not disrupted at 30°C. Moreover, PHP is possible and robust (yet imperfect) at 30°C for *GluRIIA^SP16^* mutants, but it is abolished in the case of the dominant-negative *UAS-GluRIIA^M/R^* transgene.

## DISCUSSION

We present evidence that presynaptic homeostatic potentiation (PHP) at the *Drosophila melanogaster* neuromuscular synapse is a reversible process. In doing so, we confirm prior findings showing that there is a tight inverse relationship between quantal amplitude and quantal content at the NMJ (Fig. 4). We complement those findings by conducting temperature shift experiments. We find that PHP is measurable at an early stage of larval development (Fig. 2) and can be erased over a matter of days (Figs. 3, 5). Interestingly, PHP fails or falls short of perfect compensation at high temperatures (Figs. 1, 2, 3, 6).

For the Drosophila NMJ, homeostatic potentiation is a robust and sensitive process. One assumption supported by all available data is that the larval NMJ is capable of modulating its vesicle release at any time point during development – in accordance with the presence or absence of a homeostatic challenge to synapse function. Rapid, acute induction of homeostatic signaling has previously been demonstrated at the Drosophila NMJ by application of Philanthotoxin-433 (PhTox) to impair the glutamate receptors (Frank et al., 2006), but the reversibility of this particular modulation (or any other modulation at the Drosophila NMJ) had not been studied. More generally, it is not clear what happens in metazoan nervous systems when harsh perturbations that induce homeostatic signaling are introduced for long periods of developmental time and then later removed.

### Why is reversibility slow after dominant-negative GluRIIA^M/R^ removal?

There was a robust expression of presynaptic homeostatic potentiation (PHP) for NMJs of *MHC-Gal4* >> *UAS-GluRIIA^M/R^* larvae with the *TubP-Gal80^TS^* transgene raised at 29°C for 48 hours post egg-laying (Fig. 2). Once the expression of the dominant-negative *UAS-GluRIIA^M/R^* transgene was halted, this expression of PHP was erased over a slow 48- to 72-hour period (Fig. 3). 24 hours of halted dominant-negative expression provided no relief (Fig. 3).

If PHP is a readily reversible homeostatic process, why is there a days-long time lag in order to reverse it? The answer is likely a constraint of the dominant-negative GluRIIA^M/R^ experimental perturbation, rather than a reflection of the NMJ’s capacity to respond quickly to the changed environment. In a prior study, researchers expressed functional, tagged GluRIIA trans-genic subunits at the NMJ and performed fluorescence recovery after photobleaching (FRAP) experiments (Rasse et al., 2005). Those experiments demonstrated that receptor turnover rates at the Drosophila NMJ are extremely slow: it appears that once postsynaptic densities (PSDs) reach a critical size, GluRIIA subunits are stably incorporated (Rasse et al., 2005). For our study, this likely means that the temperature downshift represented an opportunity for the NMJ to incorporate endogenous wild-type GluRIIA into a significant number of new PSDs while it continued to grow (Rasse et al., 2005; Schmid et al., 2008; Schmid et al., 2006). Given sufficient growth, the endogenously expressed GluRIIA would gradually overcome the previously incorporated dominant-negative GluRIIA^M/R^ subunits. As a result, this gradually restored neurotransmission to normal levels over a time course of 24-48 hours (Fig. 3), during which time many of the dominant-negative GluRIIA^M/R^ subunits that had previously been stably incorporated at the NMJ likely remained at the synapse.

### Reversibility of rapid and sustained forms of homeostatic plasticity

The majority of recent studies about synaptic homeostasis at the Drosophila NMJ have emphasized that presynaptic adjustments to neurotransmitter release properties must occur within minutes of drug-induced (PhTox) postsynaptic receptor inhibition in order to induce a rapid and offsetting response to PhTox challenge. Important presynaptic parameters uncovered through these studies include regulation of presynaptic Ca^2+^ influx (Frank et al., 2006; Frank et al., 2009; Müller and Davis, 2012; Wang et al., 2014; Wang et al., 2016a; Younger et al., 2013); regulation of the size of the readily releasable pool (RRP) of presynaptic vesicles (Harris et al., 2015; Müller et al., 2015; Müller et al., 2012; Wang et al., 2016a; Weyhersmüller et al., 2011); control of SNARE-mediated fusion events (Dickman and Davis, 2009; Dickman et al., 2012; Müller et al., 2011); control of neuronal excitability (Bergquist et al., 2010; Parrish et al., 2014; Younger et al., 2013); and recently, ER calcium-sensing downstream of presynaptic calcium influx (Genç et al., 2017). For almost all of the cases in which a mutation or an experimental condition blocks the short-term induction of homeostatic signaling, the same perturbation has also proven to block its long-term maintenance. Interestingly, the converse is not true. Additional studies have amassed evidence that the long-term consolidation of homeostatic signaling at the NMJ can be genetically uncoupled from its induction, and select molecules seem to be dedicated to a maintenance program that involves protein translation and signaling processes in both the neuron and the muscle (Brusich et al., 2015; Frank et al., 2009; Kauwe et al., 2016; Marie et al., 2010; Penney et al., 2012; Spring et al., 2016). These long-term processes seem to take six hours or more to take full effect (Kauwe et al., 2016).

As more molecular detail about HSP is elucidated, it will be interesting to test if the rapid induction and sustained consolidation of PHP can be reversed by similar or separate mechanisms – and what the time courses of those reversal mechanisms are. At the mouse NMJ, reversibility was recently demonstrated pharmacologically. D-Tubocurarine was applied to at a sub-blocking concentration in order to impair postsynaptic acetylcholine receptors. Within seconds of drug application, QC increased – and then within seconds of drug washout, it decreased again (Wang et al., 2016b). Follow-up experiments suggested that those rapid, dynamic changes in PHP dynamics at the mouse NMJ were mediated by a calcium-dependent increase in the size of the readily-releasable pool (RRP) of presynaptic vesicles (Wang et al., 2016b). Since there seem to be several similarities between the mouse NMJ and the Drosophila NMJ (Davis and Müller, 2015; Frank, 2014), it is possible that PHP at the insect NMJ can also be rapidly reversed.

Finally, it is instructive to examine mammalian synaptic preparations to study how homeostatic forms of synaptic plasticity are turned on and off. Groundbreaking work on cultured excitatory vertebrate synapses showed that in response to activity deprivation (or promotion), synapses employ scaling mechanisms by adding (or subtracting) AMPA-type glutamate receptors in order to counteract the perturbation (O’Brien et al., 1998; Turrigiano et al., 1998). Bidirectional scaling suggested that reversible mechanisms likely dictate homeostatic scaling processes. Complementary studies testing scaling reversibility have borne out this prediction (Rutherford et al., 1997; Swanwick et al., 2006; Wang et al., 2011). Additionally, evidence for reversible forms of homeostatic scaling have also been found in rodent sensory systems, such as auditory synapses after hearing deprivation (and restoration to reverse) (Whiting et al., 2009) and in the visual cortex after light deprivation (and restoration to reverse) (Goel and Lee, 2007). Collectively, the vertebrate and invertebrate studies support the notion that reversible fine-tuning is an efficient strategy used to stabilize activities in metazoan nervous systems. One advantage offered by the Drosophila system is a toolkit to uncover possible reversibility factors.

### Homeostatic signaling can crash at extreme temperatures

Are there environmental limitations for homeostatic potentiation at the Drosophila NMJ? Our data suggest that high temperatures represent a potential limitation on the system. It is not clear what the molecular or anatomical basis of this limit is. We do know that it is not an issue of NMJ excitation at high temperatures. This is because evoked neurotransmission for WT (or driver control) NMJs remains remarkably robust over a range of temperatures, including 30°C (Table 1). Nor does it seem to be an elimination of PHP in general because PHP was still present in the case of *GluRIIA^SP16^* animals raised at 30°C (Fig. 6, Table 1). Rather, the limitation seems to be on homeostatic signaling that supports PHP at high temperatures in the face of the dominant-negative transgene expression.

Temperature effects on neurophysiology are well documented. Recent work in crustaceans demonstrates that robust and reliable circuits like the neurons driving the rhythmicity stomatogastric nervous system can “crash” under extreme temperature challenges (Marder et al., 2015; Rinberg et al., 2013; Tang et al., 2012). For the Drosophila NMJ, prior studies of larval development documented a significant enhancement of synaptic arborization when larvae were raised at high temperatures (Sigrist et al., 2003; Zhong and Wu, 2004). Additional studies have shown that NMJ growth plasticity can be additionally affected by mutations that affect neuronal excitability (Budnik et al., 1990; Lee and Wu, 2010; Zhong et al., 1992). Given the backdrop of these data, it is not unreasonable to hypothesize that the tolerable limits of synaptic activity challenge could be different at different temperatures.

For our experiments, 29-30°C represents a potential “crash” point for homeostatic potentiation at the Drosophila NMJ. We must note, however, that our data suggest that the coping capacity of the NMJ is dependent on genotype. WT NMJs cope at all temperatures. By contrast, for dominant-negative GluRIIA-expressing NMJs, 29°C is a point at which PHP becomes imperfect (Figs. 1-2), and 30°C is a point at which it crashes (Fig. 6). For *GluRIIA^SP16^* subunit deletion NMJs, there is robust, but imperfect PHP at 30°C (Fig. 6) – not unlike the compensation seen for the dominant-negatives at 29°C. Why do these differences persist? The answer could relate to the well-documented temperature-induced alterations in NMJ growth – or alternatively, a limited availability of synaptic factors that are needed in order to cope with a double challenge of high temperature and particular impairment glutamate receptor function. Future molecular and physiological work will be needed to unravel those possibilities in the contexts of different genetic backgrounds and culturing conditions.

## ACKNOWLEDGEMENTS

We thank members of the Tootle, Lin, Wallrath, and Geyer labs for helpful discussions. Funding supporting this work includes a Whitehall Foundation Grant (2014-08-03), an NSF Grant (1557792), and an NIH/NINDS Grant (R01NS085164) to C.A.F. C.J.Y. was supported by a post-comprehensive pre-doctoral summer fellowship by the Graduate College at the University of Iowa (UI) and a Ballard and Seashore Dissertation fellowship via the Neuroscience Graduate Program at the UI.

## REFERENCES

Bergquist, S., Dickman, D.K., and Davis, G.W. (2010). A hierarchy of cell intrinsic and target-derived homeostatic signaling. Neuron 66, 220-234.

Brusich, D.J., Spring, A.M., and Frank, C.A. (2015). A single-cross, RNA interference-based genetic tool for examining the long-term maintenance of homeostatic plasticity. Frontiers in cellular neuroscience 9, 107.

Budnik, V., Koh, Y.H., Guan, B., Hartmann, B., Hough, C., Woods, D., and Gorczyca, M. (1996). Regulation of synapse structure and function by the Drosophila tumor suppressor gene dlg. Neuron 17, 627-640.

Budnik, V., Zhong, Y., and Wu, C.F. (1990). Morphological plasticity of motor axons in Drosophila mutants with altered excitability. J Neurosci 10, 3754-3768.

Burrone, J., O’Byrne, M., and Murthy, V.N. (2002). Multiple forms of synaptic plasticity triggered by selective suppression of activity in individual neurons. Nature 420, 414-418.

Corrigan, L., Redhai, S., Leiblich, A., Fan, S.J., Perera, S.M., Patel, R., Gandy, C., Wainwright, S.M., Morris, J.F., Hamdy, F., et al. (2014). BMP-regulated exosomes from Drosophila male reproductive glands reprogram female behavior. J Cell Biol 206, 671-688.

Cull-Candy, S.G., Miledi, R., Trautmann, A., and Uchitel, O.D. (1980). On the release of transmitter at normal, myasthenia gravis and myasthenic syndrome affected human end-plates. J Physiol 299, 621-638.

Daniels, R.W., Collins, C.A., Chen, K., Gelfand, M.V., Featherstone, D.E., and DiAntonio, A. (2006). A single vesicular glutamate transporter is sufficient to fill a synaptic vesicle. Neuron 49, 11-16.

Daniels, R.W., Collins, C.A., Gelfand, M.V., Dant, J., Brooks, E.S., Krantz, D.E., and DiAntonio, A. (2004). Increased expression of the Drosophila vesicular glutamate transporter leads to excess glutamate release and a compensatory decrease in quantal content. J Neurosci 24, 10466-10474.

Davis, G.W., and Bezprozvanny, I. (2001). Maintaining the stability of neural function: a homeostatic hypothesis. Annu Rev Physiol 63, 847-869.

Davis, G.W., DiAntonio, A., Petersen, S.A., and Goodman, C.S. (1998). Postsynaptic PKA controls quantal size and reveals a retrograde signal that regulates presynaptic transmitter release in Drosophila. Neuron 20, 305-315.

Davis, G.W., and Goodman, C.S. (1998). Synapse-specific control of synaptic efficacy at the terminals of a single neuron. Nature 392, 82-86.

Davis, G.W., and Müller, M. (2015). Homeostatic control of presynaptic neurotransmitter release. Annu Rev Physiol 77, 251-270.

DiAntonio, A., Petersen, S.A., Heckmann, M., and Goodman, C.S. (1999). Glutamate receptor expression regulates quantal size and quantal content at the Drosophila neuromuscular junction. J Neurosci 19, 3023-3032.

Dickman, D.K., and Davis, G.W. (2009). The schizophrenia susceptibility gene dysbindin controls synaptic homeostasis. Science 326, 1127-1130.

Dickman, D.K., Tong, A., and Davis, G.W. (2012). Snapin is critical for presynaptic homeostatic plasticity. J Neurosci 32, 8716-8724.

Featherstone, D.E., Rushton, E., Rohrbough, J., Liebl, F., Karr, J., Sheng, Q., Rodesch, C.K., and Broadie, K. (2005). An essential Drosophila glutamate receptor subunit that functions in both central neuropil and neuromuscular junction. J Neurosci 25, 3199-3208.

Frank, C.A. (2014). Homeostatic plasticity at the Drosophila neuromuscular junction. Neu ropharmacology 78, 63-74.

Frank, C.A., Kennedy, M.J., Goold, C.P., Marek, K.W., and Davis, G.W. (2006). Mechanisms underlying the rapid induction and sustained expression of synaptic homeostasis. Neuron 52, 663-677.

Frank, C.A., Pielage, J., and Davis, G.W. (2009). A presynaptic homeostatic signaling system composed of the Eph receptor, ephexin, Cdc42, and CaV2.1 calcium channels. Neuron 61, 556-569.

Gaviño, M.A., Ford, K.J., Archila, S., and Davis, G.W. (2015). Homeostatic synaptic depression is achieved through a regulated decrease in presynaptic calcium channel abundance. eLife 4.

Genç, Ö., Dickman, D.K., Ma, W., Tong, A., Fetter, R.D., and Davis, G.W. (2017). MCTP is an ER-resident calcium sensor that stabilizes synaptic transmission and homeostatic plasticity. eLife 6.

Goel, A., and Lee, H.K. (2007). Persistence of experience-induced homeostatic synaptic plasticity through adulthood in superficial layers of mouse visual cortex. J Neurosci 27, 6692-6700.

Harris, N., Braiser, D.J., Dickman, D.K., Fetter, R.D., Tong, A., and Davis, G.W. (2015). The Innate Immune Receptor PGRP-LC Controls Presynaptic Homeostatic Plasticity. Neuron 88, 1157-1164.

Hazelrigg, T., Levis, R., and Rubin, G.M. (1984). Transformation of white locus DNA in drosophila: dosage compensation, zeste interaction, and position effects. Cell 36, 469-481.

Kauwe, G., Tsurudome, K., Penney, J., Mori, M., Gray, L., Calderon, M.R., Elazouzzi, F., Chicoine, N., Sonenberg, N., and Haghighi, A.P. (2016). Acute Fasting Regulates Retrograde Synaptic Enhancement through a 4E-BP-Dependent Mechanism. Neuron 92, 1204-1212.

Kim, Y.J., Bao, H., Bonanno, L., Zhang, B., and Serpe, M. (2012). Drosophila Neto is essential for clustering glutamate receptors at the neuromuscular junction. Genes Dev 26, 974-987.

Kim, Y.J., Igiesuorobo, O., Ramos, C.I., Bao, H., Zhang, B., and Serpe, M. (2015). Prodomain removal enables neto to stabilize glutamate receptors at the Drosophila neuromuscular junction. PLoS genetics 11, e1004988.

Lee, J., and Wu, C.F. (2010). Orchestration of stepwise synaptic growth by K+ and Ca2+ channels in Drosophila. J Neurosci 30, 15821-15833.

Lnenicka, G.A., and Mellon, D., Jr. (1983a). Changes in electrical properties and quantal current during growth of identified muscle fibres in the crayfish. J Physiol 345, 261-284.

Lnenicka, G.A., and Mellon, D., Jr. (1983b). Transmitter release during normal and altered growth of identified muscle fibres in the crayfish. J Physiol 345, 285-296.

Marder, E., and Bucher, D. (2007). Understanding circuit dynamics using the stomatogastric nervous system of lobsters and crabs. Annu Rev Physiol 69, 291-316.

Marder, E., and Goaillard, J.M. (2006). Variability, compensation and homeostasis in neuron and network function. Nat Rev Neurosci 7, 563-574.

Marder, E., Haddad, S.A., Goeritz, M.L., Rosenbaum, P., and Kispersky, T. (2015). How can motor systems retain performance over a wide temperature range? Lessons from the crustacean stomatogastric nervous system. J Comp Physiol A Neuroethol Sens Neural Behav Physiol 201, 851-856.

Marder, E., and Prinz, A.A. (2002). Modeling stability in neuron and network function: the role of activity in homeostasis. Bioessays 24, 1145-1154.

Marie, B., Pym, E., Bergquist, S., and Davis, G.W. (2010). Synaptic homeostasis is consolidated by the cell fate gene gooseberry, a Drosophila pax3/7 homolog. J Neurosci 30, 8071-8082.

McGuire, S.E., Le, P.T., Osborn, A.J., Matsumoto, K., and Davis, R.L. (2003). Spatiotemporal rescue of memory dysfunction in Drosophila. Science 302, 1765-1768.

Müller, M., and Davis, G.W. (2012). Transsynaptic control of presynaptic Ca(2)(+) influx achieves homeostatic potentiation of neurotransmitter release. Curr Biol 22, 1102-1108.

Müller, M., Genç, Ö., and Davis, G.W. (2015). RIM-Binding Protein Links Synaptic Homeostasis to the Stabilization and Replenishment of High Release Probability Vesicles. Neuron 85, 1056-1069.

Müller, M., Liu, K.S., Sigrist, S.J., and Davis, G.W. (2012). RIM Controls Homeostatic Plasticity through Modulation of the Readily-Releasable Vesicle Pool. J Neurosci 32, 16574-16585.

Müller, M., Pym, E.C., Tong, A., and Davis, G.W. (2011). Rab3-GAP controls the progression of synaptic homeostasis at a late stage of vesicle release. Neuron 69, 749-762.

Murthy, V.N., Schikorski, T., Stevens, C.F., and Zhu, Y. (2001). Inactivity produces increases in neurotransmitter release and synapse size. Neuron 32, 673-682.

O’Brien, R.J., Kamboj, S., Ehlers, M.D., Rosen, K.R., Fischbach, G.D., and Huganir, R.L. (1998). Activity-dependent modulation of synaptic AMPA receptor accumulation. Neuron 21, 1067-1078.

Parrish, J.Z., Kim, C.C., Tang, L., Bergquist, S., Wang, T., Derisi, J.L., Jan, L.Y., Jan, Y.N., and Davis, G.W. (2014). Kruppel mediates the selective rebalancing of ion channel expression. Neuron 82, 537-544.

Penney, J., Tsurudome, K., Liao, E.H., Elazzouzi, F., Livingstone, M., Gonzalez, M., Sonenberg, N., and Haghighi, A.P. (2012). TOR is required for the retrograde regulation of synaptic homeostasis at the Drosophila neuromuscular junction. Neuron 74, 166-178.

Petersen, S.A., Fetter, R.D., Noordermeer, J.N., Goodman, C.S., and DiAntonio, A. (1997). Genetic analysis of glutamate receptors in Drosophila reveals a retrograde signal regulating presynaptic transmitter release. Neuron 19, 1237-1248.

Plomp, J.J., Van Kempen, G.T., De Baets, M.B., Graus, Y.M., Kuks, J.B., and Molenaar, P.C. (1995). Acetylcholine release in myasthenia gravis: regulation at single end-plate level. Ann Neurol 37, 627-636.

Plomp, J.J., van Kempen, G.T., and Molenaar, P.C. (1992). Adaptation of quantal content to decreased postsynaptic sensitivity at single endplates in alpha-bungarotoxin-treated rats. J Physiol 458, 487-499.

Ramos, C.I., Igiesuorobo, O., Wang, Q., and Serpe, M. (2015). Neto-mediated intracellular interactions shape postsynaptic composition at the Drosophila neuromuscular junction. PLoS genetics 11, e1005191.

Rasse, T.M., Fouquet, W., Schmid, A., Kittel, R.J., Mertel, S., Sigrist, C.B., Schmidt, M., Guzman, A., Merino, C., Qin, G., et al. (2005). Glutamate receptor dynamics organizing synapse formation in vivo. Nat Neurosci 8, 898-905.

Redhai, S., Hellberg, J.E., Wainwright, M., Perera, S.W., Castellanos, F., Kroeger, B., Gandy, C., Leiblich, A., Corrigan, L., Hilton, T., et al. (2016). Regulation of Dense-Core Granule Replenishment by Autocrine BMP Signalling in Drosophila Secondary Cells. PLoS genetics 12, e1006366.

Rinberg, A., Taylor, A.L., and Marder, E. (2013). The effects of temperature on the stability of a neuronal oscillator. PLoS Comput Biol 9, e1002857.

Rongo, C., and Kaplan, J.M. (1999). CaMKII regulates the density of central glutamatergic synapses in vivo. Nature 402, 195-199.

Rutherford, L.C., DeWan, A., Lauer, H.M., and Turrigiano, G.G. (1997). Brain-derived neurotrophic factor mediates the activity-dependent regulation of inhibition in neocortical cultures. J Neurosci 17, 4527-4535.

Schmid, A., Hallermann, S., Kittel, R.J., Khorramshahi, O., Frolich, A.M., Quentin, C., Rasse, T.M., Mertel, S., Heckmann, M., and Sigrist, S.J. (2008). Activity-dependent site-specific changes of glutamate receptor composition in vivo. Nat Neurosci 11, 659-666.

Schmid, A., Qin, G., Wichmann, C., Kittel, R.J., Mertel, S., Fouquet, W., Schmidt, M., Heckmann, M., and Sigrist, S.J. (2006). Non-NMDA-type glutamate receptors are essential for maturation but not for initial assembly of synapses at Drosophila neuromuscular junctions. J Neurosci 26, 11267-11277.

Schuster, C.M., Davis, G.W., Fetter, R.D., and Goodman, C.S. (1996a). Genetic dissection of structural and functional components of synaptic plasticity. I. Fasciclin II controls synaptic stabilization and growth. Neuron 17, 641-654.

Schuster, C.M., Davis, G.W., Fetter, R.D., and Goodman, C.S. (1996b). Genetic dissection of structural and functional components of synaptic plasticity. II. Fasciclin II controls presynaptic structural plasticity. Neuron 17, 655-667.

Sigrist, S.J., Reiff, D.F., Thiel, P.R., Steinert, J.R., and Schuster, C.M. (2003). Experience-dependent strengthening of Drosophila neuromuscular junctions. J Neurosci 23, 6546-6556.

Spring, A.M., Brusich, D.J., and Frank, C.A. (2016). C-terminal Src Kinase Gates Homeostatic Synaptic Plasticity and Regulates Fasciclin II Expression at the Drosophila Neuromuscular Junction. PLoS genetics 12, e1005886.

Staley, B.K., and Irvine, K.D. (2010). Warts and Yorkie mediate intestinal regeneration by influencing stem cell proliferation. Curr Biol 20, 1580-1587.

Swanwick, C.C., Murthy, N.R., and Kapur, J. (2006). Activity-dependent scaling of GABAergic synapse strength is regulated by brain-derived neurotrophic factor. Mol Cell Neurosci 31, 481-492.

Tang, L.S., Taylor, A.L., Rinberg, A., and Marder, E. (2012). Robustness of a rhythmic circuit to short- and long-term temperature changes. J Neurosci 32, 10075-10085.

Thiagarajan, T.C., Lindskog, M., and Tsien, R.W. (2005). Adaptation to synaptic inactivity in hippocampal neurons. Neuron 47, 725-737.

Turrigiano, G.G. (2008). The self-tuning neuron: synaptic scaling of excitatory synapses. Cell 135, 422-435.

Turrigiano, G.G. (2017). The dialectic of Hebb and homeostasis. Philos Trans R Soc Lond B Biol Sci 372.

Turrigiano, G.G., Leslie, K.R., Desai, N.S., Rutherford, L.C., and Nelson, S.B. (1998). Activity-dependent scaling of quantal amplitude in neocortical neurons. Nature 391, 892-896.

Ueda, A., and Wu, C.F. (2015). The role of cAMP in synaptic homeostasis in response to environmental temperature challenges and hyperexcitability mutations. Frontiers in cellular neuroscience 9, 10.

Wang, H.L., Zhang, Z., Hintze, M., and Chen, L. (2011). Decrease in calcium concentration triggers neuronal retinoic acid synthesis during homeostatic synaptic plasticity. J Neurosci 31, 17764-17771.

Wang, T., Hauswirth, A.G., Tong, A., Dickman, D.K., and Davis, G.W. (2014). Endostatin Is a Trans-Synaptic Signal for Homeostatic Synaptic Plasticity. Neuron 83, 616-629.

Wang, T., Jones, R.T., Whippen, J.M., and Davis, G.W. (2016a). alpha2delta-3 Is Required for Rapid Transsynaptic Homeostatic Signaling. Cell reports 16, 2875-2888.

Wang, X., Pinter, M.J., and Rich, M.M. (2016b). Reversible Recruitment of a Homeostatic Reserve Pool of Synaptic Vesicles Underlies Rapid Homeostatic Plasticity of Quantal Content. J Neurosci 36, 828-836.

Wefelmeyer, W., Puhl, C.J., and Burrone, J. (2016). Homeostatic Plasticity of Subcellular Neuronal Structures: From Inputs to Outputs. Trends Neurosci 39, 656-667.

Weyhersmüller, A., Hallermann, S., Wagner, N., and Eilers, J. (2011). Rapid active zone remodeling during synaptic plasticity. J Neurosci 31, 6041-6052.

Whiting, B., Moiseff, A., and Rubio, M.E. (2009). Cochlear nucleus neurons redistribute synaptic AMPA and glycine receptors in response to monaural conductive hearing loss. Neuroscience 163, 1264-1276.

Wierenga, C.J., Ibata, K., and Turrigiano, G.G. (2005). Postsynaptic expression of homeostatic plasticity at neocortical synapses. J Neurosci 25, 2895-2905.

Younger, M.A., Müller, M., Tong, A., Pym, E.C., and Davis, G.W. (2013). A presynaptic ENaC channel drives homeostatic plasticity. Neuron 79, 1183-1196.

Zhong, Y., Budnik, V., and Wu, C.F. (1992). Synaptic plasticity in Drosophila memory and hyperexcitable mutants: role of cAMP cascade. J Neurosci 12, 644-651.

Zhong, Y., and Wu, C.F. (2004). Neuronal activity and adenylyl cyclase in environment-dependent plasticity of axonal outgrowth in Drosophila. J Neurosci 24, 1439-1445.

